# Rare Gli1+ perivascular fibroblasts promote skin wound repair

**DOI:** 10.1101/2022.05.16.491785

**Authors:** Xiaoyan Sun, Karl Annusver, Tim Dalessandri, Maria Kasper

## Abstract

Growing evidence suggests that perivascular cells play important roles in tissue repair of various organs. In the skin, the contribution and importance of these cells for wound repair is not resolved. Here we demonstrate that a specific Gli1+ subpopulation residing in the perivascular niche serves as an important cellular source for wound healing fibroblast. First, we show that *Gli1* expression marks small subsets of both pericytes and perivascular adventitial cells. Upon injury both cell types rapidly responded already within their original niche, however only the progeny of Gli1+ adventitial cells expanded and differentiated into wound-contracting myofibroblasts. Genetic ablation of these cells significantly impaired wound healing, which was associated with the reduction of aSMA+ myofibroblast-mediated wound contraction. After wound closure these cells reverted to an aSMA-negative fibroblast state, and intriguingly, they persisted in wounds over long term and adopted a non-fibrotic fibroblast signature. In sum, our data sheds new light on the functional diversity of perivascular-cell subtypes in the skin, and proposes a new mesenchymal cell source that promotes wound healing.

## INTRODCUTION

Cutaneous wound healing is a highly orchestrated process that involves distinct but interrelated phases: hemostasis, inflammation, proliferation, and remodeling ^1^. Each of these phases is dependent on the active involvement of a variety of epithelial, mesenchymal and immune cell types ^2^. In recent years, evidence is mounting that mesenchymal cells within the perivasculature niche contribute to more than just vessel formation and stability during wound repair (reviewed in ^3–5^). For example, subsets of pericytes and perivascular fibroblasts can give rise to wound-induced myofibroblasts and be involved in fibrotic scar formation upon injury in a wide variety of organs such as kidney, muscle, lung, brain, spinal cord, and heart ^4,6–9^. However, whether or not pericytes (situated in the capillaries’ tunica intima) and adventitial fibroblasts (embedded in the tunica adventitia of bigger vessels) have differential functions despite both sharing a vascular niche, is not adequately resolved. In particular, the role of the perivascular adventitial fibroblasts in skin wound healing remains largely unexplored due to the lack of mouse models for cell-type specific fate mapping.

Recent studies in various organs other than the skin have shown that *Gli1*-expression marks subsets of the perivascular network, including pericytes and adventitial fibroblasts ^10–12^. This prompted us to test whether *Gli1*-expression can be utilized to investigate the role of pericytes and/or adventitial fibroblasts in skin wound repair. As most of the body parts in adult mammalian skin are at any given time in *telogen* (no active hair growth) ^13^, we systematically characterized *Gli1*-expression in adult mouse telogen skin, which revealed a small number of Gli1+ cells residing in the perivascular niches of small and large vessels. Using lineage tracing in combination with full-thickness wounding, targeted cell ablation, and quantitative image analysis, we reveal the differential dynamics, contribution, and the fate of Gli1+ pericytes and adventitial fibroblasts in short- and long-term wound repair, and the consequence of their absence.

## RESULTS

### Identification of Gli1+ cells in nerve and vascular niches of telogen mouse skin

The long resting phase of hair growth called *2^nd^ telogen* starts in postnatal week 6-7 and can last up to week 12. We utilized this long period of defined (static) tissue architecture as it allowed us to precisely determine the placement of Gli1+ cells prior to wounding, and during all the early critical wound repair phases. In telogen skin, *Gli1* has been shown to mark specific epithelial and mesenchymal cell populations in association with the hair follicle and touch dome ^14^. Additionally, throughout the dermis and subcutaneous fat, we and others have observed rare and scattered, yet undefined Gli1+ cells ^14,15^, which we hypothesized to inhabit the perivascular niche.

To characterize these rare cells, we first analyzed the back skin of 8-week-old Gli1^lacZ^ reporter mice, where *Gli1*-expressing cells are visualized via b-galactosidase staining (blue cells). Tissue histology shows that Gli1+ cells are represented in all previously described structures (Figure 1a), and the scattered cells primarily reside in the perivascular and nerve niches of the hypodermis, muscles, and deep facia (Figure 1b). For in-depth characterization of these cells, we crossed Gli1-CreERT2 with Rosa26-tdTomato mice (hereafter called Gli1^TOM^) to enable inducible genetic labeling (i.e.: lineage tracing) of Gli1+ cells. Lineage tracing was induced via administration of tamoxifen at 8 weeks of age, and the dorsal skin was collected one week later (Figure 1c). Immunostaining against a panel of markers revealed that some Gli1^TOM^ cells were interspersed in the NFH+ nerve bundles that were characterized as Schwann cells (SOX10+ PDGFRβ–) (Figure 1c, lower panel). However, Gli1^TOM^ cells were also present in the perivascular niche constituting a subpopulation of both pericytes as well as adventitial fibroblasts. The Gli1^TOM^ pericytes (PDGFRβ+ NG2+) were intimately associated with the microvasculature in direct contact to CD31-expressing endothelial cells and fully surrounded by a Collagen IV+ basement membrane (Figure 1c, mid panel), and the Gli1^TOM^ adventitial fibroblasts (PDGFRβ+ aSMA–) were localized in the adventitia of larger vessels, adjacent to the aSMA+ smooth muscle layer and distant from CD31+ endothelial cells (Figure 1c, upper panel; Figure S1).

**Figure 1.**
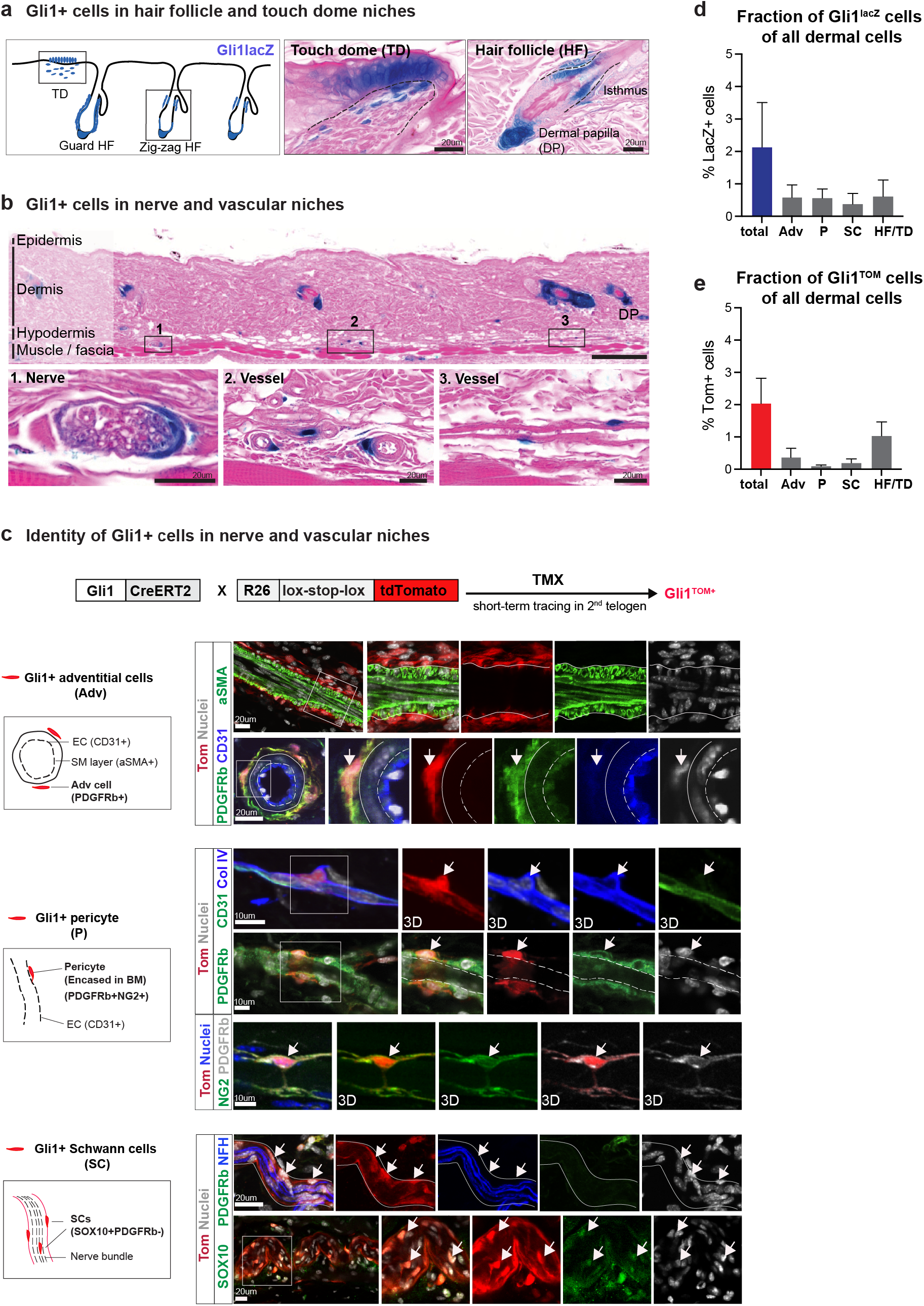
Identification of Gli1+ cells in nerve and vascular niches. **(a-b)** X-galactosidase staining on back skin of Gli1^lacZ^ mice at 2^nd^ telogen stage. (a) Illustration and representative images of LacZ-positive cells in HF/TD niches. Dashed lines indicate epidermal-dermal border. (b) A panorama of dorsal skin with magnified areas showing LacZ-positive cells in the nerve and vascular niches. **(c)** Generation of Gli1creERT2;Rosa26-tdTomato (hereafter: Gli1^TOM^) mice. Mice at 2^nd^ telogen stage were treated with tamoxifen to enable lineage-tracing of Gli1+ cells. Gli1^TOM^ skin was stained against various markers as indicated for cell type identification in nerve and vascular niches (right panel). Arrows denote Gli1^TOM^ cells expressing respective markers corresponding to left panel cartoon and lines indicate the nerve or vessel niches. Boxes indicate the magnified areas. 3D represents 3D image visualization from depth-blended views from any angle. BM, base membrane; EC, endothelial cells; SM, smooth muscle. **(d)** Quantification of total and each fraction of LacZ+ cells of all dermal cells in Gli1^lacZ^ skin (6 big panoramas from n=3 mice). Data are shown as means ± SEM. **(e)** Quantification of total and each fraction of Tom+ cells of all dermal cells in Gli1^TOM^ skin (6 big panoramas from n=3 mice). Data are shown as means ± SEM. Adv, adventitial cells; P, pericytes; SC, Schwann cells; HF, hair follicles; TD, touch dome; DP, dermal papilla. Scale bars: 200um (b panorama); 20um (a, b upper and lower panel, c); 10um (b mid panel).

### Fate tracing of the Gli1+ cells in healed wounds

As *Gli1* expression marked overall a very small subset of dermal cells (around 2%) with perivascular cells accounting for 0.5-1% (Figure 1d-e), we first wanted to know if these Gli1+ cells would indeed generate a notable wound bed contribution. Therefore, we treated Gli1^TOM^ mice with tamoxifen at 8 weeks, created a full thickness wound (4 mm punch) in the back skin after 5 days, and analyzed the wounds 12 days later when the wound re-epithelialization was just completed (Figure 2a). Indeed, in addition to the expected epithelial cell contribution (Figure 2b) ^14^, we observed numerous dermal Gli1^TOM^ cells streaming from the surrounding tissue into the wound, thereby contributing a considerable fraction of cells to the wound bed (Figure 2c-d). Notably, *Gli1* expression itself is entirely absent in short- and long-term wounds (Figure S2a-d) ^16^, suggesting that the *Gli1*-traced cells in wounds may even adopt a different identity than in healthy skin.

**Figure 2.**
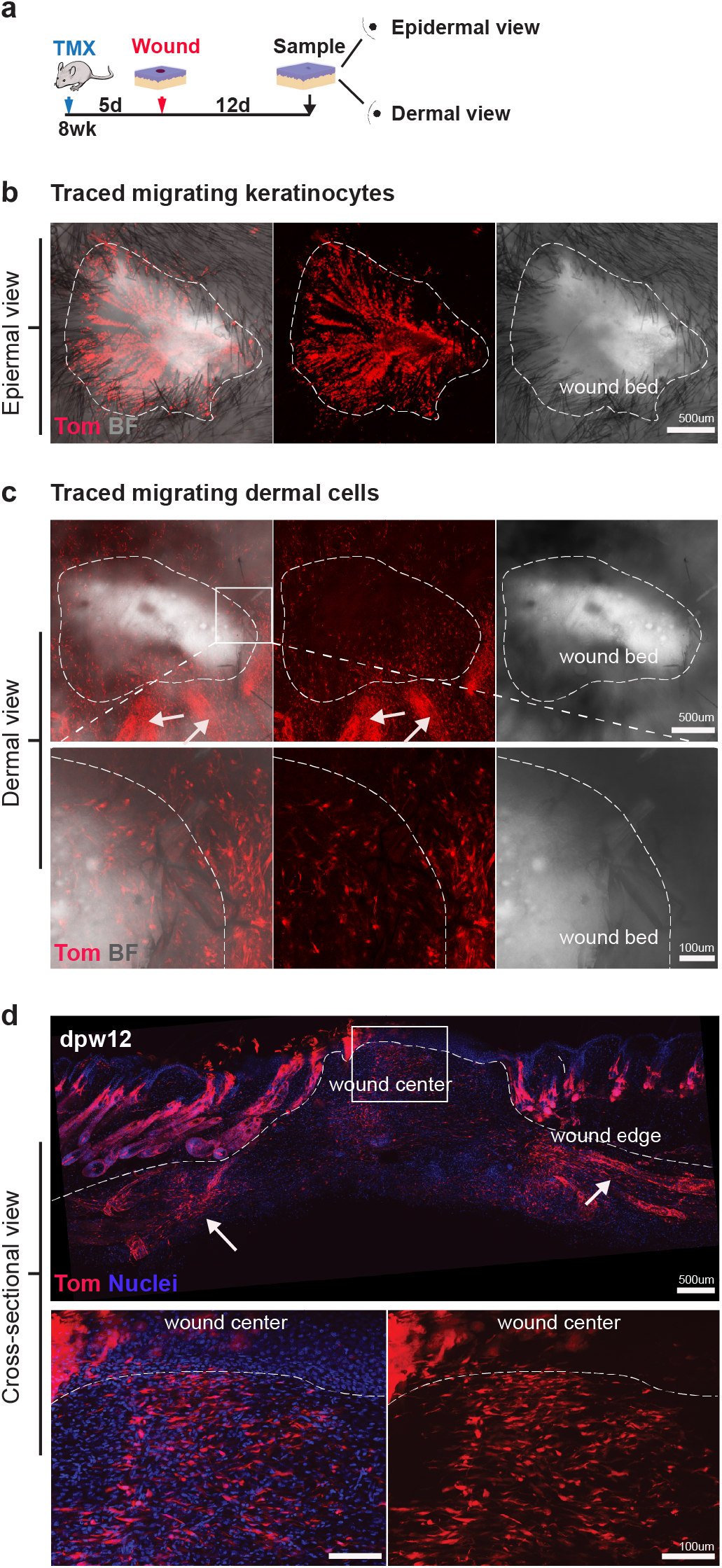
Contribution of Gli1+ cells to wound bed upon full-thickness wounding. **(a)** Experimental timeline. Briefly, Gli1^TOM^ mice at 8 weeks of age were treated with tamoxifen. 5 days later, full-thickness wounds were created on the dorsal skin and sampled at 12 days post wounding (dpw) (n=25 mice). **(b)** Epidermal view of a flat-mounted wound. **(c)** Dermal view of a flat-mounted wound. Arrows indicate the migrating streams of Gli1^TOM^ cells in wound surrounding area. **(d)** A cross-sectional view of a horizonal whole mount wound and a magnified region of wound center. Arrows indicate the Gli1^TOM^ cells in wound edge. Dashed line marks the epidermal-dermal border and wound margin. Scale bars: 500um (b, c-d panorama); 100um (c-d insets).

To characterize the cell types that the Gli1^TOM^ cells represent in the wound bed, and if these cells remain the same linages as in healthy skin, we first stained the wounds with SOX10 (pan-glial) and Laminin (encasing pericytes), in combination with CD31 (endothelium) and aSMA (vascular smooth muscle). As we would have expected, some of the Gli1^TOM^ cells became again glial cell (Figure 3a,e) or pericyte lineage (Figure 3b,c,e). Of note, we were unable to spot if Gli1^TOM^ cells also became again adventitial fibroblasts due to the very sparce presence of aSMA+ vessels in wound neodermis (Figure S3a-b). However, the majority of Gli1^TOM^ cells in the wound bed were neither glial cells nor pericytes or adventitial cells, and via CD45 staining we also ruled out unexpected Gli1^TOM^ immune cell contribution (Figure 3f). Next, we examined the wound bed to see if these Gli1^TOM^ cells would become normal wound fibroblasts marked by the pan-fibroblast marker PDGFRa (Figure 3g). Intriguingly, most of the Gli1^TOM^ cells in the wound bed were indeed fibroblasts (Figure 3e), demonstrating that Gli1+ cells mainly contribute to the fibroblast pool in response to wounding.

**Figure 3.**
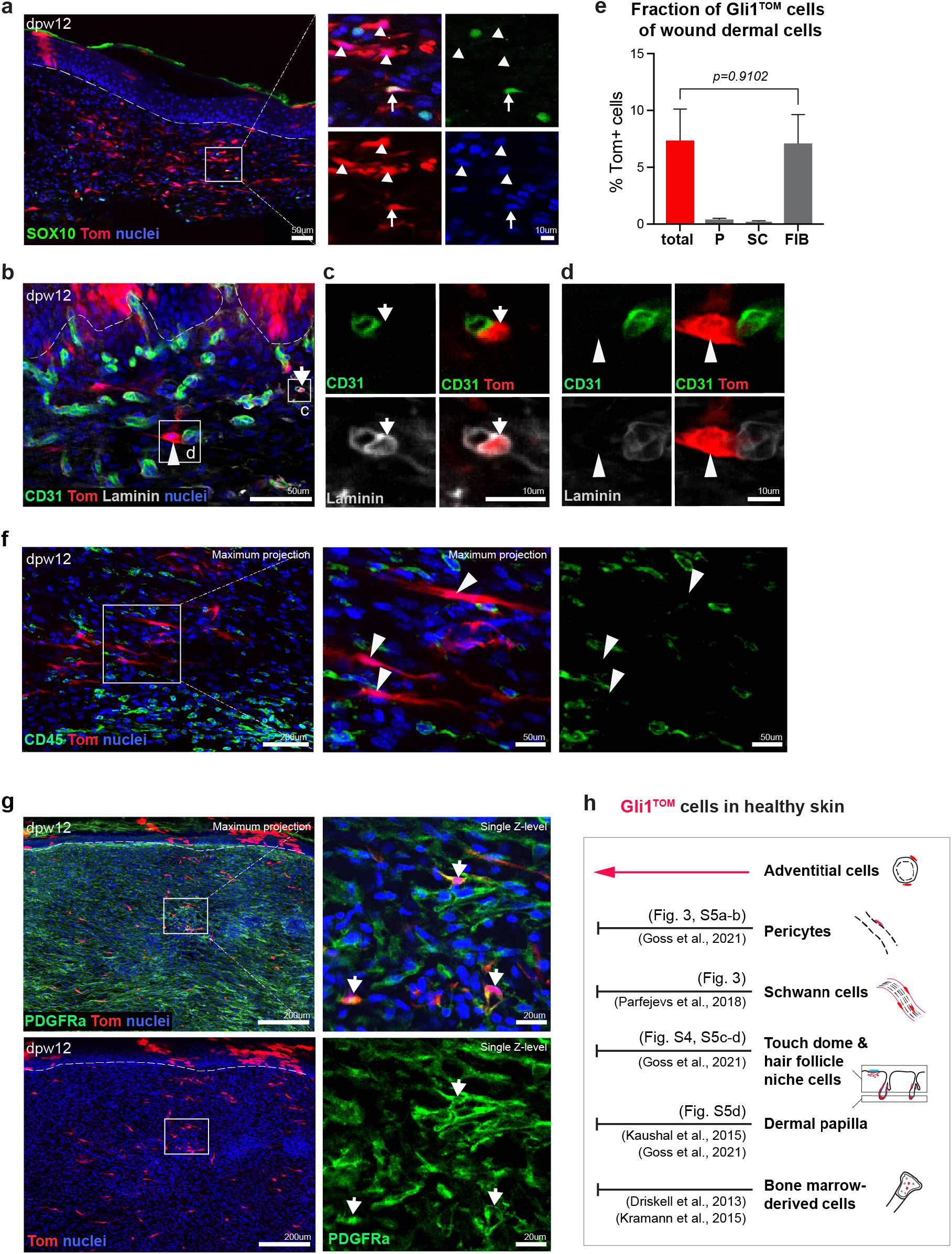
Fate tracing of Gli1+ cells in wound neo-dermis. **(a)** Immunostaining for SOX10 (pan-glial marker) in dpw12 wounds. Arrows indicate SOX10-positive Gli1^TOM^ cells while arrowheads SOX10-negative Gli1^TOM^ cells. **(b)** Immunostaining for CD31 (endothelium marker) and pan-Laminin (encased pericytes) in dpw12 wounds. **(c-d)** Magnified areas of Gli1^TOM^ cells sitting close to endothelium. Arrows and arrowheads indicate a Gli1^TOM^ cell covered with (c) or without (d) Laminin, respectively. **(e)** Quantification of total and each fraction of Gli1^TOM^ cells in wound neo-dermis at 12dpw (n=3 mice). P, pericytes; SC, Schwann cells; FIB, fibroblasts. Data are shown as means ± SEM. Unpaired two-tailed t-test. *p* value of comparisons between total Gli1^TOM^ cells and fibroblast fraction is shown in the graph. **(f)** Immunostaining for CD45 (myeloid cell marker) in dpw12 wounds. A magnified area showing CD45-negative Gli1^TOM^ cells (arrowheads). **(g)** Immunostaining for PDGFRa (pan-fibroblast marker) in dpw12 wounds. A magnified area showing PDGFRa-positive Gli1^TOM^ cells (arrows). **(h)** Illustration summarizing all possible wound-contributing cell types from Gli1^TOM^ cells in healthy skin. Scale bars: 200um (f-g panorama); 50um (a-b panorama, f-g insets); 10um (a-b insets).

### Adventitial fibroblasts are the major source of the Gli1^TOM^ traced wound neo-dermis

Next, we investigated the possible cell source for Gli1^TOM^ fibroblasts in the wound bed. To pinpoint the cells of origin, we systematically assessed dermal wound contribution of all possible *Gli1*-expressing cell types (i.e.: hair follicle and touch dome associated fibroblasts, dermal papilla cells, pericytes, adventitial fibroblasts, glial/Schwann cells, and bone-marrow derived cells) (summarized in Figure 3h).

Based on previous studies, bone marrow derived cells ^10,17^ as well as dermal papilla cells ^18,19^ could convincingly be excluded as a potential source. Moreover, as Sox10 is a well-established marker for cells of glial origin incl. Schwann cells ^20^, and glial/Schwann cells are lineage restricted during skin wound repair ^21^, we also could exclude this cell lineage as the cell source for the Gli1^TOM^ traced fibroblast population in the wound neo-dermis.

To probe the hair follicle and touch dome niche fibroblasts as a cellular origin, we made use of the fact that *Gli1* expression in the hair follicle and touch dome niches (shown in Figure 1a) depends on the presence of cutaneous nerve signals ^14^. Utilizing a cutaneous denervation protocol in Gli1^LacZ^ mice, we more specifically found that *Gli1*-expression is not only lost in the nerve-ending cells of the hair follicle isthmus and touch dome but also in the nearby fibroblasts, while it remains in the large nerve bundles and perivascular niche (Figure S4a). Next, we used Gli1^TOM^ mice where we first abolished *Gli1*-expression in hair follicle and touch dome fibroblasts (via denervation), followed by labeling of the remaining *Gli1*-expressing cells (via tamoxifen administration) and subsequent wounding 5 days later (Figure S4b). The wound-introducing biopsies were used to confirm successful denervation (i.e.: lack of Tomato tracing hair follicle isthmus and touch dome niche cells) (Figure S4c-e). Twelve days post wounding, we quantified the dermal Gli1^TOM^ cell contribution in both the denervated as well as sham-operated (control) wound sites, revealing some but not significant decrease (Figure S4f-g). This indicates that Gli1+ fibroblasts of the hair follicle and touch dome niches do not serve as a major cellular source of the wound bed.

Lastly, recent work of Goss and colleagues ^19^ – combined with our new results in Gli1^TOM^ mice – allowed us to exclude the Gli1+ pericytes as a cell source for the wound bed contribution. They used NG2^Cre^R26^TOM^ mice which labelled more than 70-80% of all pericytes in telogen skin. NG2^TOM^ traced cells contributed only to the pericyte lineage during wound repair ^19^, and crucially, we found that all Gli1+ pericytes co-express NG2 in healthy skin (Figure S5a-b). Thus, Gli1^TOM^ pericytes cannot give rise to the common wound fibroblast pool. Interestingly, our Gli1^TOM^-traced cells in the touch dome and hair follicle (incl. dermal papilla) also express NG2, which further supports that those cells do not make a major contribution to the wound bed (Figure S5c-d). However, importantly, the Gli1+ adventitial fibroblasts in healthy skin do not express NG2 (Figure S6).

Thus, we can conclude that the adventitial fibroblasts constitute the major Gli1+ cell source for dermal wound repair (Figure 3e,g). Strikingly, this contribution is coming from a minor fraction of *Gli1*-expressing cells in healthy skin (Gli1^LacZ^: 0.58%; Gli1^TOM^: 0.37%, Figure 1d-e) that significantly expands to contribute to the wound neo-dermis fibroblast pool.

### Gli1+ adventitial fibroblasts become wound healing myofibroblasts

To reveal the initial behavior of the *Gli1*-expressing adventitial fibroblasts in response to full-thickness wounding, we analyzed 2^nd^-telogen-labeled Gli1^TOM^ cells in wounds 2- and 6-days post wounding (2dpw and 6dpw). Two hours prior to sacrifice, a single pulse of bromodeoxyuridine (BrdU) was given to label the actively proliferating cells (Figure 4a).

**Figure 4.**
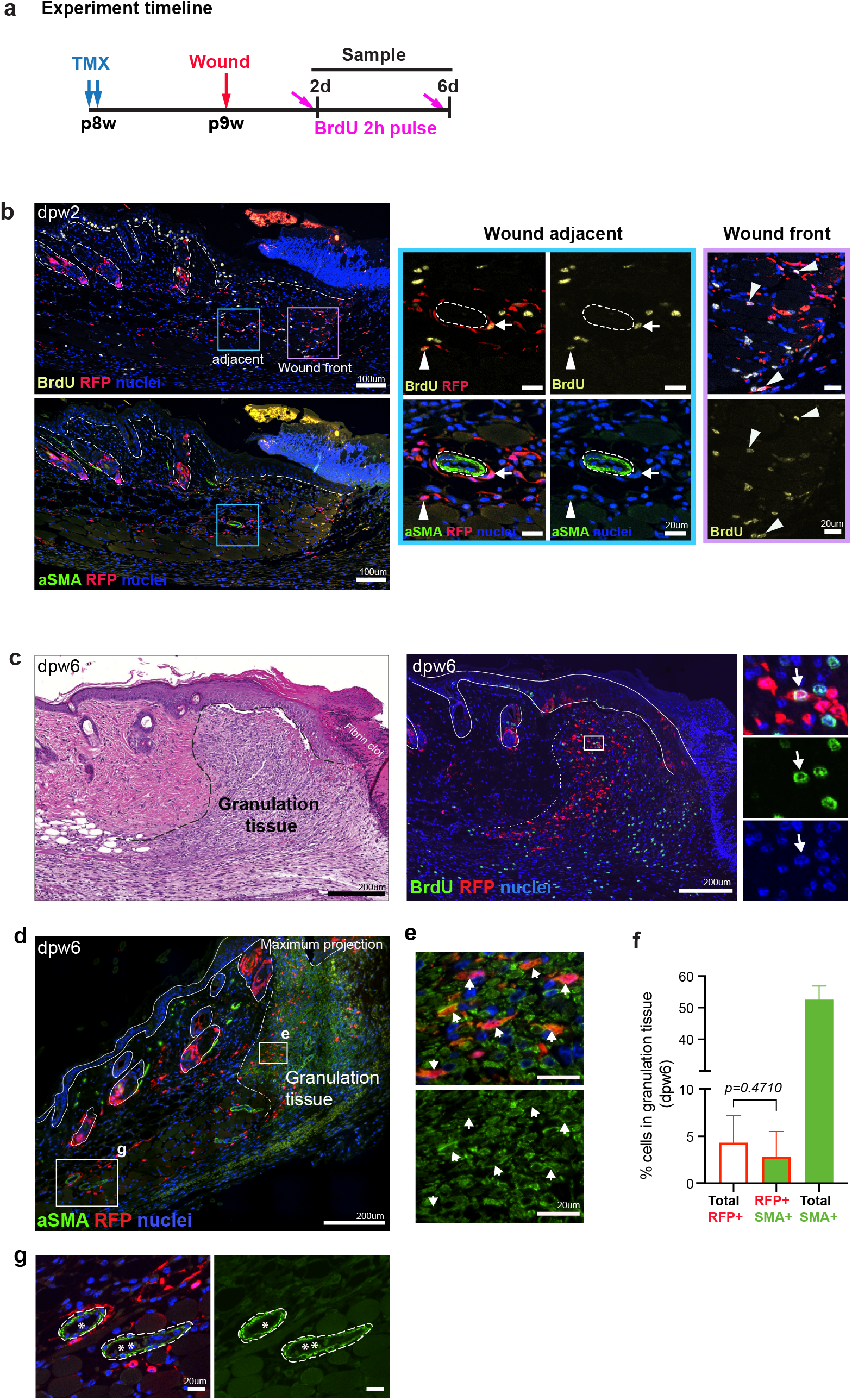
Expansion and differentiation of Gli1+ cells upon wounding. **(a)** Experimental timeline. Briefly, wounds were sampled at 2 or 6 days post wounding (dpw) and single pulse of BrdU was given i.p. 2 hours prior to sampling (n=3 mice each). **(b)** Immunostaining for BrdU, aSMA and RFP (Tomato) in dpw2 wounds. Magnified areas of wound adjacent and wound front are shown. Arrow indicates a BrdU-positive Gli1^TOM^ cell remaining in the vessel adventitia, while arrowheads indicate the scattered BrdU-positive Gli1^TOM^ cells. (**c**) Immunostaining for BrdU and RFP (Tomato) in dpw6 wounds, with an H&E-stained serial section showing the equivalent wound area. Arrows indicate BrdU-positive Gli1^TOM^ cells. **(d)** Immunostaining of aSMA and RFP (Tomato) in dpw6 wounds. Lines indicate skin epidermal-dermal border and the wound margin. **(e)** Magnified area of wound front from (d). Arrows indicate the double positive cells. **(f)** Quantification of RFP+ and aSMA+ cells in granulation tissue of dpw6 wounds (n=3 mice). Data are shown as means ± SEM. Unpaired two-tailed t-test. *p* value of comparisons between total RFP+ cells and its aSMA+ fraction is shown in the graph. **(g)** Magnified area of wound adjacent area from (d). Asterisks indicate aSMA+ vessels. Dashed lines indicate the relative location of RFP+ cells surrounding aSMA+ layer. Scale bars: 200um (c panorama and d); 100um (b panaroma); 20um (b and d insets and g).

During skin homeostasis, *Gli1*-expressing cells were mostly non-proliferative (Figure S7). However, upon full-thickness wounding, Gli1^TOM^ cells underwent rapid activation and expansion. At 2dpw, Gli1^TOM^ adventitial fibroblasts were already proliferating before leaving their vessel niche, as well as expanded at the wound front outside their original niche (Figure 4b). At 6dpw – a timepoint during highly active granulation tissue formation – numerous Gli1^TOM^ cells were present in the granulation tissue with some of the cells still proliferating (Figure 4c).

As myofibroblasts are characteristic for wound granulation tissue, we used the well-known myofibroblast marker alpha-smooth muscle actin (aSMA) to determine if Gli1^TOM^ cells would contribute to the myofibroblast pool. Indeed, at 6dpw, the majority of all Gli1^TOM^ cells in wound granulation tissue were aSMA+ (Figure 4d-f). Reassuringly, Gli1^TOM^ cells adjacent to the wounds remained negative for aSMA expression (shown is an example of adventitious fibroblasts; Figure 4g).

We conclude that Gli1+ adventitial fibroblasts quickly respond to acute wounding, starting to proliferate already within their original niche. Subsequently they leave their niche while further expanding within the wound bed and change their identity to wound-contracting myofibroblasts.

### Genetic ablation of Gli1+ cells results in delayed wound healing

To unveil the functional importance of the Gli1+ cell contribution for wound healing, we utilized inducible diphtheria toxin receptor mice (Rosa26-iDTR) crossed with Gli1-CreERT2 mice (hereafter called Gli1^iDTR^). Upon tamoxifen treatment, expression of the human DTR in *Gli1*-expressing cells allows for the specific ablation of the *Gli1*-lineage via diphtheria toxin administration (Figure 5a).

**Figure 5.**
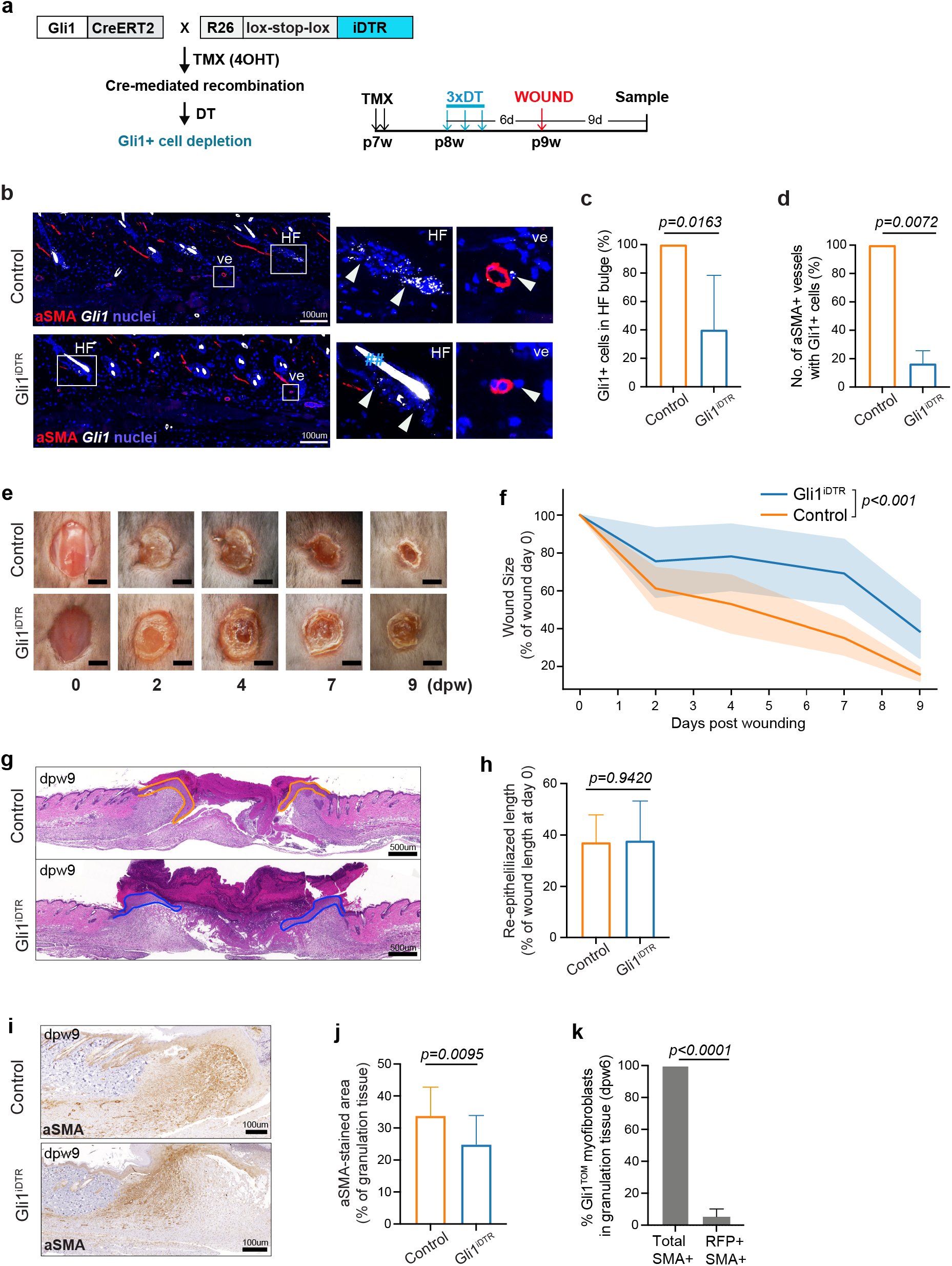
Depletion of Gli1+ cells delays wound repair. **(a)** Generation of Gli1-CreERT2;Rosa26-iDTR (hereafter: Gli1^iDTR^) mice and the experimental timeline. Briefly, mice at 7 weeks of age were treated with 4-hydroxyl tamoxifen (TMX). One week later, mice were treated with diphtheria toxin (DT) for 3 times with one day interval. 6 days after 1^st^ DT treatment, full-thickness wounds were created on the dorsal skin and sampled at 9 days post wounding (dpw). **(b)** *Gli1* mRNA and aSMA antibody co-staining on skin biopsies of both genotypes 6 days after DT treatment. Magnified areas showing a hair follicle and an aSMA+ vessel from each genotype. ## indicates autofluorescence of hair shaft. **(c-d)** Quantification of *Gli1+* cells in hair follicle bulge (c) and the number of aSMA+ vessels accompanied with *Gli1+* cells (d). Data are shown as percentage of the control mice with means ± SEM. n=2 mice each. Unpaired two-tailed t-test. **(e)** Representative digital photos of wounds from both genotypes at different healing time points. **(f)** Wound closure (size) of both genotypes at different healing time points (n=5 each). Data are shown as proportional to initial wound size at dpw0. Statistics was done using ANOVA for linear models (see Methods). *P* value is shown in the graph. **(g)** Representative images of wounds at dpw9. Solid line indicates new epithelial tongue. **(h)** The length of new epithelial tongue was measured. Data is shown as percentage of the wound length at day 0 with means ± SEM (n=5 each). Unpaired two-tailed t-test. **(i)** Representative images of aSMA staining at dpw9 wounds. **(j)** Quantification of aSMA-stained area at dpw9 wounds. Data are shown as percentage of granulation tissue area (n=5 mice each) with means ± SEM. Unpaired two-tailed t-test. **(k)** Quantification of total aSMA+ myofibroblasts and RFP+ myofibroblasts at dpw6 wounds in Gli1^TOM^ mice (n=3). Data are shown as means ± SEM. Unpaired two-tailed t-test. Scale bars: 2mm (e); 100um (b and i); 500um (g).

Notably, in our trial experiments we observed a clear delay of wound healing even in wild type mice treated with diphtheria toxin in comparison with those without diphtheria toxin treatment (Figure S9a). Therefore, to control for the effect of diphtheria toxin on wound healing, we chose an experimental setup where we treated both Gli1^iDTR^ (Gli1creERT2;iDTR^homo^) and control littermates (iDTR^homo^ or iDTR^het^) with diphtheria toxin (Figure 5a). In brief, we treated control and Gli1^iDTR^ mice with tamoxifen at 7 weeks (activating DTR expression in Gli1+ cells), activated DTR-mediated cell ablation at 8 weeks (via diphtheria toxin administration), created a full thickness wound in the back skin at 9 weeks, and analyzed the wounds 9 days later (Figure 5a).

The ablation efficiency of DTR-expressing cells was assessed at the time of wounding and collection (6 and 15 days after the first diphtheria toxin treatment, respectively). Based on qRT-PCR on whole skin lysates, the DTR expression in Gli1^iDTR^ mice was reduced by 78,3% (at 6 days) and 90% (at 15 days) compared to the controls (Figure S9b). Importantly, *Gli1* mRNA staining demonstrated significant reduction of *Gli1*-expressing cells in both the epidermal (e.g.: hair follicle bulge) and dermal compartments (e.g.: adventitial fibroblasts) at the time of wounding, suggesting a sufficient depletion of Gli1+ cells in Gli1^iDTR^ mice (Figure 5b-d).

Indeed, analysis of the wound size over the healing timeline (0, 2, 4, 7, 9 days after wounding) revealed a significantly slower overall healing process in Gli1^iDTR^ mice compared to the control mice (*p*<0.001; see Methods) (Figure 5e-f). The wound-closure delay in Gli1^iDTR^ was already visible at day 4 and became significant at days 7 and 9 post wounding (Figure 5e,f). Remarkably, at day 9 the wound area in Gli1^iDTR^ mice was in average 2.4 times larger than in control mice (Figure 5f; see Methods). As diphtheria toxin cell ablation in Gli1^iDTR^ mice also targets some epithelial cell compartments (Figure 1a) ^14^, we ruled out that this wound-healing delay did not stem from a slower re-epithelialization process (Figure 5g-h). As the majority of *Gli1*-progeny in wound granulation tissue became myofibroblasts (see Figure 4f), we examined if the delay of wound closure comes from a reduction of wound myofibroblasts upon Gli1+ cell ablation. Indeed, elimination of Gli1+ cells significantly reduced the aSMA+ myofibroblast pool in 9-day-old granulation tissue signified by a 21% decrease of aSMA-stained areas (Figure 5i-j). This decrease is striking as Gli1^TOM^ cells represent only 5.4% of all myofibroblasts (Figure 5k).

Taken together, these results implicate that depletion of Gli1+ cells results in slower wound repair, which is primarily associated with a reduced number of myofibroblasts that originate from Gli1+ adventitial fibroblasts.

### Gli1^TOM^ myofibroblasts return to a normal dermal fibroblast phenotype and persist in the healed dermis after long-term remodeling

When analyzing 2- and 6-months-old wounds of Gli1^TOM^ mice we observed a substantial persistent tracing of fibroblasts in the healed dermis (Figure 6a-c). Normally, when a wound is closed myofibroblasts disappear via apoptosis or accelerated senescence ^8,22,23^, and only under certain conditions (e.g.: regression of liver fibrosis) myofibroblasts were observed to revert to long-term wound fibroblasts ^24^. Thus, it was intriguing to investigate the identity of the long-term contributing Gli1^TOM^ fibroblasts in more detail. During active wound repair (12d) Gli1^TOM^ (myo)fibroblasts were spread out over the entire wound bed (Figure 2c), however over time (2-6 months) they preferentially occupy the lower part of the wound bed (Figure 6b-c). We have previously described 3 distinct fibroblast subtypes in healthy mouse skin, with one subtype displaying two distinct states during hair cycling: *Dcn*^high^ (upper dermis, telogen), *Sparc*^high^ (upper dermis, anagen), *Gpx3*^high^ (lower dermis above panniculus carnosus muscle (PCM)), and *Plac8*^high^ (below PCM) ^25^. Interestingly, in 6-month-old wounds *Dcn*^high^ and *Sparc*^high^ cells are almost mutually exclusive (Figure 6c). They divide the wound dermis into an upper (*Sparc*^high^) and a lower (*Dcn*^high^) part; thus, the majority of Gli1^TOM^ cells are *Dcn*^high^ and *Sparc*^low/-^ (Figure 6c-d). Occasionally, we also found *Gpx3* expression in some of these cells (Figure 6c-d).

**Figure 6.**
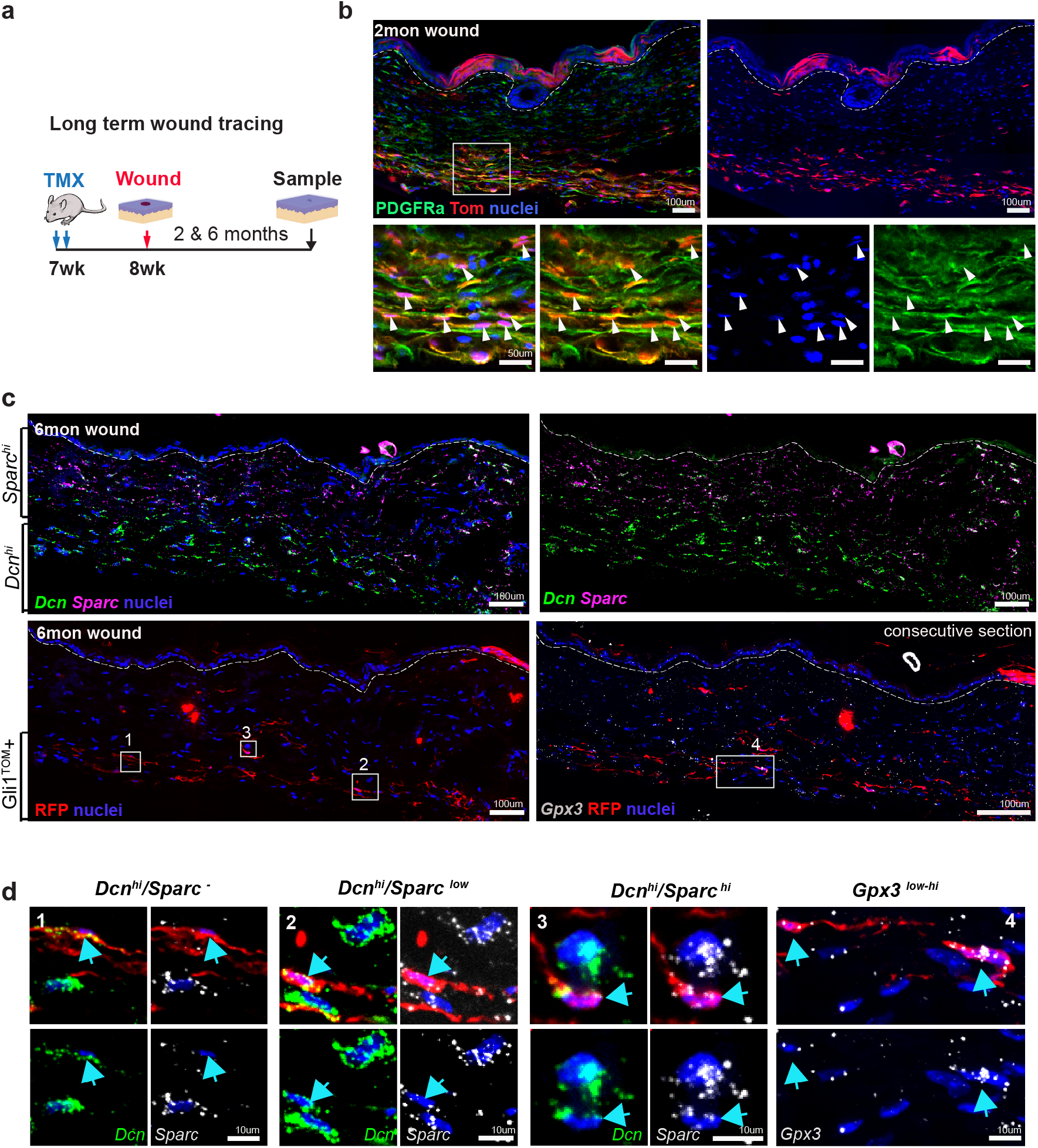
Identity of Gli1^TOM^ fibroblasts in long-term wound. **(a)** Experimental scheme. Briefly, full-thickness wounds were collected after 2 and 6 months post wounding (n=3 mice). **(b)** Immunostaining for PDGFRa (pan-fibroblast marker) in 2-month-old wounds. Box shows a magnified area of wound lower region. Arrowheads indicate the PDGFRa-positive Gli1^TOM^ cells. Dashed lines indicate the epidermal-dermal border. **(c)** 6-month-old wounds were stained for single-molecule RNA-FISH probes of different fibroblast lineage markers *Dcn, Sparc* and *Gpx3* in combination with RFP antibody. Dashed lines indicate the epidermal-dermal border. **(d)** Magnified areas of wound Gli1^TOM^ fibroblasts with different markers (arrows). Scale bars: 100um (b and c); 10um (d).

In sum, the majority (if not all) of the *Gli1*-progeny of adventitial fibroblasts contribute to the myofibroblast pool during wound closure, after which they return to an aSMA-negative wound fibroblast state persisting long-term and taking on a *Sparc*-negative *Dcn*-positive fibroblast identity. The latter is particularly interesting as *Sparc* encodes a matricellular protein (i.e., a dynamically expressed glycoprotein modulating extracellular matrix interactions) that is commonly overexpressed in fibrotic diseases ^26^, which may potentially endow Gli1^TOM^ cells with an anti-fibrotic role in cutaneous wound healing.

## DISCUSSION

*Gli1*-expressing cells have been studied in various organs other than the skin and found to represent a network of perivascular cells with mesenchymal stem cells properties ^10–12^. However, in the skin, the identity, dynamics, and fate of Gli1+ cells as well as their functional role upon full-thickness wounding have not yet been explored. In this study, we systematically characterized *Gli1*-expressing cells in mouse telogen skin and revealed that a specific Gli1+ subpopulation in the perivascular niche – namely the Gli1+ adventitial fibroblasts – serves as an important cellular source for wound healing fibroblasts.

The myofibroblast composition and source can differ between organs, injury models and size of the original wound ^27–29^. In the skin, for example, many cell types including local dermal fibroblasts ^17,30^, fascia cells ^31^, dermal adipocyte-derived cells ^32^ and hematopoietic-lineage cells ^33^ can generate myofibroblasts in the wound context. Earlier studies in kidney, muscle, lung, brain, spinal cord and heart have demonstrated that perivascular cells including pericytes can also contribute to the myofibroblast pool upon organ injury ^4,6–9^. However, more recent studies in some of the same (heart, brain) and additional (fat, skin and skeletal muscle) organs have demonstrated that pericytes only give rise to pericytes and do not function as progenitors for fibroblasts, although pericytes do proliferate upon injury (Figure S8) ^19,34,35^. This discrepancy is likely attributed to several factors such as the type of organs, severity of injury models, a non-unified nomenclature for the different perivascular cell subtypes (adventitial fibroblasts, pericytes, or potential pericyte subtypes), as well as lack of a comprehensive knowledge about Cre-driver mouse lines and cell-type specific protein expression. Here in this work, we mapped all Gli1+ cell populations *in situ* (Gli1^LacZ^ and short-term tracing of Gli1^TOM^) to differentiate pericytes and adventitial fibroblasts in order to unveil their respective role upon full-thickness wounding. Intriguingly, (at least) in the skin, it’s the adventitial fibroblasts, but not the pericytes, that contribute to the myofibroblast pool and thereby play key roles in the healing of wounds. Since the wound healing process in most mammalian organs are remarkably similar irrespective of the underlying etiology ^2^, our findings in skin wound healing can be informative to wound repair more generally.

Our genetic cell ablation experiment shows that the depletion of Gli1+ cells prior to wounding caused a significant reduction of the aSMA+ myofibroblast pool, which we predominantly attribute to the ablation of Gli1+ adventitial fibroblasts. However, as *Gli1* is also expressed in small subsets of Schwann cells and pericytes, it is possible that the simultaneous ablation of these minor Gli1+ populations (see Figure 3e) contributes to the wound healing delay even though these cell types do not become myofibroblasts themselves ^19,21,35^. It has been reported that peripheral glial cells can support tissue repair by paracrine signaling e.g. regulating the number of myofibroblasts in the skin ^21,36^, while Gli1+ pericytes were found essential for a certain type of vessel formation during bone repair ^37^.

Lastly, an unexpected and intriguing finding of this work was that after wound closure Gli1+ adventitial fibroblast progeny eventually returned to a normal fibroblast phenotype, persisting in wounds over long-term and acquiring a non-fibrotic fibroblast signature. Myofibroblasts usually disappear via apoptosis or senescence when the acute wound has closed ^8,22,23^, and the persistence of myofibroblasts is often associated with pathological conditions such as hypertrophic scarring or undesired scar contracture ^38,39^. Thus, this work opens for new explorations to investigate the mechanism of myofibroblast inactivation or reversion into a normal quiescent-state fibroblast, as this may identify new approaches to revert or reduce hypertrophic scar formation.

## METHODS

### Mice

All transgenic mouse lines in this study were obtained from Jackson Laboratory, and bred on a C57BL/6J background in-house under pathogen-free conditions. Details of the mice are listed in Key Resource Table. Adult mice in the second telogen stage of the hair cycle (7-11 weeks old) were used and all procedures were performed on dorsal skin. For each condition and time point, at least three mice were treated and analyzed, unless stated otherwise. The experimental mice were housed at the animal facility of Karolinska Institute. All animal experiments were carried out in accordance with Swedish legislation and approved by the Stockholm South or Linköping Animal Ethics Committees.

### Lineage tracing

For genetic lineage tracing, the Gli1CreERT2 mice were crossed with Rosa26-tdTomato mice (hereafter: Gli1^TOM^). To induce Cre recombinase activity, the estrogen analog tamoxifen (20mg/ml dissolved in corn oil) was given by intraperitoneal (i.p.) injection at a dose of 3 mg for twice or at single dose of 4 or 6 mg prior to wounding. A mix of males and females mice were used in the experiments and similar results were obtained. For BrdU incorporation, mice were injected i.p. with BrdU (Sigma) solution (10 mg/ml) at a dose of 0.1 mg/g body weight 2 hours prior to sacrifice.

### Gli1+ cell ablation

For ablation of Gli1+ cells, Gli1creERT2 mice were crossed with Rosa26-iDTR mice (hereafter: Gli1^iDTR^). Littermates without Gli1CreERT2 were used as control (hereafter: Control). All mice were treated topically with 2 doses of 0.75 mg 4-Hydroxytamoxifen (4-OHT, dissolved in 100ul acetone) to induce Cre recombination. Diphtheria toxin (DT) was then injected i.p. at a dose of 40ng/g body weight every other day for total 3 times before wounding. Skin biopsies were collected at the time of wounding as well as 6 or 15 days after DT treatments for ablation efficiency analysis. For consistency and better comparison, the control and experimental mice were all sex- and age-matched.

### Full-thickness excisional wounding

For fate tracing, full-thickness excisional wounds were created by 4 mm punch (Acuderm) on the dorsal skin of mice. The wounds were left uncovered for mimicking the natural healing process. The day when the wounds were made was considered as day 0. For tracking the behavior of lineage-traced cells, the wounded regions were harvested at different time points as stated in the figures or text. For evaluating the cell ablation effect on wound repair between control and cell ablated mice, full-thickness wounds were created by 5 mm punch (Acuderm), and closely examined daily. Wounds were photographed by digital camera for recording wound closure changes and wound size was measured afterwards by ImageJ software.

### Evaluation of wound healing parameters in cell-ablated mice

Wound re-epithelialization was assessed by measuring the length of newly formed wound epithelium on H&E-stained sections from the middle of wounds at day 9. Wound closure (changes of wound size) was measured based on macroscopic wound images taken at different time points during wound healing process, which were carried out using Image J software. In a series of cross-cut wound sections, the ones with the widest wound gap (center of wound) were chosen for further immunohistochemical staining.

### Unilateral surgical denervation of dorsal skin

A full-thickness midline incision was made on the dorsal skin and cutaneous nerves on the right back were surgically removed as previously described ^40^. The contralateral skin was left untouched and served as control. The midline incision was closed with 6.0 nylon suture afterwards. For comparative study of wound healing in control and denervated sides, 4 mm punch (Acuderm) was used for creating full-thickness excisional wounds on both sides. Wound healing process was monitored and healed wounds were collected for comparison of fate tracing between control and denervated side.

### Tissue preparation

Freshly obtained healthy or wounded skin samples were collected and processed in different ways. For gross assessment of tdTomato lineage tracing in whole wound area, whole wounds were flat-mounted between two coverslips, and scanned from both epidermal and dermal side by Nikon A1R confocal microscope. For paraffin-embedded (FFPE) samples, tissue were fixed in 4% paraformaldehyde (PFA) for 24 hrs and then embedded in paraffin. Paraffin blocks were processed into 4 um-thickness skin sections. For whole mount (HWM) samples, tissues were fixed in 4% PFA for 20 min and mounted in OCT embedding medium (Histolab). Subsequently, 60-150 um sections were cut with a cryostat. For later RNA extractions, tissue were submerged and stored in RNAlater solution (Invitrogen) immediately after harvest to inactivate RNases and stabilize RNA.

### Histological analysis and Immunohistochemistry on FFPE samples

For histological analysis, FFPE sections were directly counterstained with hematoxylin and eosin. For immunostaining, tissue sections, after de-waxing, were subjected to antigen retrieval by 10 mM citrate buffer (DAKO) or Diva Decloaker (Biocare Medical), then blocked with serum and incubated with primary antibodies (listed in Key Resource Table). Biotinylated secondary antibodies (Vector Laboratories, 1:500) or Alexa Fluor Dyes 488, 546, 647 or 680 (Invitrogen, 1:500) were used for detecting signals from primary antibodies.

### Immunostaining on HWM samples

Skin tissue was sliced into 100-150 um thick using a Cryostat. Tissue strips were blocked with PB buffer (0.1% fish skin gelatin, 0.5% Triton X-100 and 0.5% skimmed milk powder in PBS) and stained as previously described ^41^. TO-PRO-3 (Molecular Probes) or Hoechst (Invitrogen) were used for nuclear staining.

### Single molecule (sm) RNA-FISH

For characterizing Gli1-lineage traced fibroblasts in long-term wounds, smRNA-FISH of *Dcn, Sparc, Gpx3* as well as *Gli1* was performed using RNAscope Multiplex Fluorescent V2 Assay (Advanced Cell Diagnostics) using TSA with Cy3, Cy5, and/or Fluorescein (NEL760001KT, Perkin Elmer) on FFPE sections. Subsequent immuno-staining of RFP antibody and nuclear counterstaining was performed in combination with smRNA-FISH according to manufacturer’s instructions. Images were acquired on a Nikon A1R spinning disk confocal as tiled images (10%–15% stitching).

### LacZ (β-Galactosidase) staining

Fresh skin tissue was fixed in 4% PFA for 30 min at RT and then washed briefly with rinse buffer (2 mM MgCl2, 0.01% Nonidet P-40 in PBS). After 18 hrs incubation in the β-galactosidase substrate solution (1 mg/mL X-Gal, 5 mM K3Fe(CN)6, 5 mM K4Fe (CN)60·3H2O in rinse buffer) at 37 °C in the dark, the stained tissues were embedded in paraffin, then processed into 4 um tissue sections and counterstained with eosin or H&E.

### Taqman assay

RNA was extracted from skin biopsies with an RNEasy Mini Plus Kit (QIAGEN) and Tissue Lyzer II (QIAGEN), DNA-eliminated with a RNAse-free DNAse kit (QIAGEN), and then cDNA was synthesised with a SuperScript II kit (ThermoFisher). TaqMan™ qRT-PCR performed with human HBEGFR probe and mouse *Ppia* probe (housekeeping gene) with Taqman Advanced Master Mix (ThermoFisher).

### Image acquisition and processing

Imaging was performed using LSM710-NLO confocal microscope (Zeiss), a Nikon A1R confocal microscope, or Panoramic digital slide scanner (3DHistech Ltd.). Images were processed using NIS-Elements software (Nikon), Zen 2009 software (Zeiss), or Fiji (ImageJ 1.53c) software. Some acquired images were occasionally optimized for brightness, contrast, and color balance.

### Quantitative image analysis

Fluorescence images were preprocessed (maximum intensity projection, masking) with Fiji (ImageJ 1.53c) and subsequently quantified with a combination of CellProfiler (3.1.9), custom scripts using a gaussian mixture model for stained cell detection (scikit-learn 0.24.2) and manual verification of the results. Brightfield images (immunohistochemical staining with DAB) were quantified with a combination of QuPath software (v0.2.0-m9) for pixel classification and custom scripts using a gaussian mixture model for cell classification. Specifically, for better discrimination between different cell types based on markers in combination with anatomical location, manual quantification was performed for each cell type in (1) Gli1^lacZ^ healthy skin (Figure 1d), (2) Gli1^Tom^ lineage-traced healthy skin (Figure 1e), (3) Gli1^Tom^ lineage-traced wound (Figure 3e).

### Statistical analysis

Data on barplots are represented as means ± SEM from at least three independent experiments, unless stated otherwise. Statistical analyses were carried out using the Prism version 9.0 (GraphPad Software). Statistical significance was determined using an unpaired two-tailed t-test. Exact *P* values are shown in the figures. A *p* value of less than 0.05 was considered significant.

### Wound closure statistical analysis

Wound closure rate between Control and Gli1^iDTR^ mice is shown as mean and the 95% CI (Figure 5f). To analyze the statistical significance of overall healing process between two groups, we first constructed cubic linear models (statsmodels.regression.linear_model.OLS) that explain the wound closure proportion depending either only on the day (H_0_) or on a combination of the day and the genotype (H_1_) (see below). Subsequently, ANOVA (statsmodels.stats.anova.anova_lm) was used to compare if the model with the addition of genotype (H_1_) is better at explaining the observed data than the model using only days (H_0_).

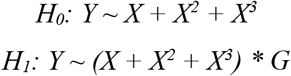

Y: proportion of wound closure, X: day post wounding, G: genotype.

## KEY RESOURCE TABLE

| REAGENT or RESOURCE | SOURCE | IDENTIFIER | Additional information |
| --- | --- | --- | --- |
| **Mouse Lines** |  |  |  |
| Gli1CreERT2 | 42 | JAX: 007913; RRID:IMSR_JAX:007913 |  |
| Gli1LacZ | 42 | JAX: 008211; RRID:IMSR_JAX:008211 |  |
| Rosa26-tdTomato | 43 | JAX: 007908; RRID:IMSR_JAX:007908 |  |
| Rosa26-iDTR | 44 | JAX: 007900; RRID:IMSR_JAX:007900 |  |
| **Antibodies** |  |  |  |
| aSMA | Abcam | Cat# ab5694; RRID:AB_2223021 | 1:200 IF;<br>1:1000 IHC |
| aSMA | Sigma-Aldrich | Cat# C6198; RRID:AB_476856 | 1:250 IF |
| aSMA | Leica Biosystems | Cat# NCL-SMA; RRID:AB_442134 | 1:50 IF |
| BrdU | Serotec | Cat# OBT0030G; RRID:AB_609567 | 1:400 IF |
| CD140b (PDGFRβ) | Biolegend | Cat# 136008; RRID:AB_2268091 | 1:100 IF |
| CD31 | BD pharmingen | Cat# 557355; RRID:AB_396660 | 1:200 IF |
| CD31 | Abcam | Cat# ab28364; RRID:AB_726362 | 1:200 IF |
| CD45 | Biolegend | Cat# 103116; RRID:AB_312981 | 1:100 IF |
| Collagen IV | Abcam | Cat# ab19808; RRID:AB_445160 | 1:50 IF |
| Laminin | Abcam | Cat# ab11575; RRID: AB_298179 | 1:200 IF |
| NFH (Neurofilament heavy chain) | Aves/MediQip | Cat# NFH; RRID:AB_2313552 | 1:1000 IF |
| PDGFRa | R&D | Cat# AF1062; RRID:AB_2236897 | (10 ug/ml) IF |
| RFP | Rockland | Cat# 600-401-379; RRID:AB_220971 | 1:200 IF |
| SOX10 (N-20) | Santa Cruz | Cat# sc-17342; RRID:AB_2195374 | 1:1000 IF |
| NG2 | Millipore | Cat# AB5320; RRID:AB_1121367 | 1:200 IF |
| TO-PRO-3 | Invitrogen | Cat# T3605 | 1:1000 |
| Hoechst 33342 | Invitrogen | Cat# H3570 | 1 ug/ml |
| **Critical Reagents and Commercial Assays** |  |  |  |
| Tamoxifen | Sigma | Cat# T5648 |  |
| 4-OH Tamoxifen | Sigma | Cat# H6278 |  |
| Diphtheria toxin | Sigma | Cat# D0564 |  |
| RNAscope Multiplex<br>Fluorescent Reagent<br>Kit v2 | ACDBio/Bio-Techne | Cat# 323100 |  |
| TSA Cy 3, Cy 5,<br>TMR, Fluorescein<br>Evaluation Kit | Perkin Elmer | Cat# NEL760001KT |  |
| **RNA probes** |  |  |  |
| *3-plex Positive*<br>*Control Probe* | ACDBio/Bio-Techne | Cat# 320881 |  |
| *3-plex Negative*<br>*Control Probe* | ACDBio/Bio-Techne | Cat# 320871 |  |
| *Mm-Gli1* | ACDBio/Bio-Techne | Cat# 311001 |  |
| *Mm-Dcn* | ACDBio/Bio-Techne | Cat# 413281 |  |
| *Mm-Sparc-C2* | ACDBio/Bio-Techne | Cat# 466781-C2 |  |
| *Mm-Gpx3-C3* | ACDBio/Bio-Techne | Cat# 509901-C3 |  |
| **Software and Algorithms** |  |  |  |
| Fiji (ImageJ 1.53c) | <sup>45,46</sup> |  |  |
| CellProfiler v3.1.9 | <sup>47</sup> |  |  |
| QuPath software<br>(v0.2.0-m9) | <sup>48</sup> |  |  |
| scikit-learn 0.24.2 | <sup>49</sup> |  |  |

## Acknowledgements

We would like to thank Maryam Saghafian for help with skin denervation surgery, Alexandra Are for technical assistance with stainings, and Igor Adameyko, Tina Jacob and Lars Jakobsson for valuable discussions and feedback during manuscript preparation. This work was supported by grants from the Wenner-Gren Foundation to T.D., Karolinska Institutet (KID) to K.A., Swedish Research Council (2018-02963), Swedish Cancer Society (CAN 2018/793), Ragnar Söderberg Foundation (M127/12), LEO Foundation (GOLD Award), Stiftelsen Gösta Miltons Donationsfond, and Karolinska Institutet (2-2111/2019) to M.K. Parts of this study were performed at the Live Cell Imaging facility/Nikon Center of Excellence, Karolinska Institutet, supported by grants from the Swedish Research Council, KI infrastructure, and Centre for Innovative Medicine.

## Author contributions

XS and MK conceptualized the study, designed the experiments, interpreted the data and wrote the manuscript. XS conducted all the animal-related work including lineage tracing, surgeries and cell ablation experiments. XS and KA performed the stainings and imaging. KA performed the image processing and quantifications with the help of XS. TD performed the qPCR analysis and assisted with the DT-depletion experiments. All authors discussed the results and provided input at all stages.

## Supplementary Information

**Figure S1.** Gli1^TOM^ tracing in the adventitia of different types of larger vessels.

(Related to Figure 1)

**Figure S2.** Absence of *Gli1* expression in healed wound.

(Related to Figure 2)

**Figure S3.** Gli1^TOM^ tracing in association with aSMA+ vessels in wound neo-dermis.

(Related to Figure 3)

**Figure S4.** Gli1^TOM^ tracing in association with hair follicle and touch dome niches.

(Related to Figure 3)

**Figure S5.** NG2 expression in pericytes, touch dome and hair follicle niches of Gli1^TOM^ skin.

(Related to Figure 3)

**Figure S6.** NG2 is not expressed in Gli1^TOM^ adventitial cells.

(Related to Figure 3)

**Figure S7.** Gli1+ cells are not actively proliferating during skin homeostasis.

(Related to Figure 4)

**Figure S8.** Upon wounding Gli1+ cells already start proliferating within their original niches.

(Related to Figure 4)

**Figure S9.** Diphtheria toxin administration affects mouse skin wound repair.

(Related to Figure 5)

## REFERENCES

1. Ellis, S., Lin, E. J. & Tartar, D. Immunology of Wound Healing. Curr Dermatology Reports 7, 350–358 (2018).

2. Gurtner, G. C., Werner, S., Barrandon, Y. & Longaker, M. T. Wound repair and regeneration. Nature 453, 314–321 (2008).

3. Mills, S. J., Cowin, A. J. & Kaur, P. Pericytes, Mesenchymal Stem Cells and the Wound Healing Process. Cells 2, 621–634 (2013).

4. Birbrair, A. et al. Type-1 pericytes accumulate after tissue injury and produce collagen in an organ-dependent manner. Stem Cell Res Ther 5, 122 (2014).

5. Carlo, S. E. D. & Peduto, L. The perivascular origin of pathological fibroblasts. J Clin Invest 128, 54–63 (2018).

6. Göritz, C. et al. A Pericyte Origin of Spinal Cord Scar Tissue. Science 333, 238–242 (2011).

7. Humphreys, B. D. et al. Fate Tracing Reveals the Pericyte and Not Epithelial Origin of Myofibroblasts in Kidney Fibrosis. Am J Pathology 176, 85–97 (2010).

8. Dulauroy, S., Carlo, S. E. D., Langa, F., Eberl, G. & Peduto, L. Lineage tracing and genetic ablation of ADAM12(+) perivascular cells identify a major source of profibrotic cells during acute tissue injury. Nat Med 18, 1262–1270 (2012).

9. Lin, S.-L., Kisseleva, T., Brenner, D. A. & Duffield, J. S. Pericytes and perivascular fibroblasts are the primary source of collagen-producing cells in obstructive fibrosis of the kidney. Am J Pathology 173, 1617–1627 (2008).

10. Kramann, R. et al. Perivascular Gli1+ Progenitors Are Key Contributors to Injury-Induced Organ Fibrosis. Cell Stem Cell 16, 51–66 (2015).

11. Schneider, R. K. et al. Gli1+ Mesenchymal Stromal Cells Are a Key Driver of Bone Marrow Fibrosis and an Important Cellular Therapeutic Target. Cell Stem Cell 20, 785–800.e8 (2017).

12. Kramann, R. et al. Adventitial MSC-like Cells Are Progenitors of Vascular Smooth Muscle Cells and Drive Vascular Calcification in Chronic Kidney Disease. Cell Stem Cell 19, 628–642 (2016).

13. Geyfman, M., Plikus, M. V., Treffeisen, E., Andersen, B. & Paus, R. Resting no more: re-defining telogen, the maintenance stage of the hair growth cycle. Biol Rev 90, 1179–1196 (2015).

14. Brownell, I., Guevara, E., Bai, C. B., Loomis, C. A. & Joyner, A. L. Nerve-Derived Sonic Hedgehog Defines a Niche for Hair Follicle Stem Cells Capable of Becoming Epidermal Stem Cells. Cell Stem Cell 8, 552–565 (2011).

15. Sun, X. et al. Coordinated hedgehog signaling induces new hair follicles in adult skin. Elife 9, e46756 (2020).

16. Lim, C. H. et al. Hedgehog stimulates hair follicle neogenesis by creating inductive dermis during murine skin wound healing. Nat Commun 9, 4903 (2018).

17. Driskell, R. R. et al. Distinct fibroblast lineages determine dermal architecture in skin development and repair. Nature 504, 277–281 (2013).

18. Kaushal, G. S. et al. Fate of Prominin-1 Expressing Dermal Papilla Cells during Homeostasis, Wound Healing and Wnt Activation. J Invest Dermatol 135, 2926–2934 (2015).

19. Goss, G., Rognoni, E., Salameti, V. & Watt, F. M. Distinct Fibroblast Lineages Give Rise to NG2+ Pericyte Populations in Mouse Skin Development and Repair. Frontiers Cell Dev Biology 9, 675080 (2021).

20. Britsch, S. et al. The transcription factor Sox10 is a key regulator of peripheral glial development. Gene Dev 15, 66–78 (2001).

21. Parfejevs, V. et al. Injury-activated glial cells promote wound healing of the adult skin in mice. Nat Commun 9, 236 (2018).

22. Jun, J.-I. & Lau, L. F. The Matricellular Protein CCN1/CYR61 Induces Fibroblast Senescence and Restricts Fibrosis in Cutaneous Wound Healing. Nat Cell Biol 12, 676–685 (2010).

23. Desmoulière, A., Redard, M., Darby, I. & Gabbiani, G. Apoptosis mediates the decrease in cellularity during the transition between granulation tissue and scar. Am J Pathology 146, 56–66 (1995).

24. Kisseleva, T. et al. Myofibroblasts revert to an inactive phenotype during regression of liver fibrosis. Proc National Acad Sci 109, 9448–9453 (2012).

25. Joost, S. et al. The Molecular Anatomy of Mouse Skin during Hair Growth and Rest. Cell Stem Cell 26, 441–457.e7 (2020).

26. Wang, J.-C. et al. Attenuation of fibrosis in vitro and in vivo with SPARC siRNA. Arthritis Res Ther 12, R60–R60 (2010).

27. Micallef, L. et al. The myofibroblast, multiple origins for major roles in normal and pathological tissue repair. Fibrogenesis Tissue Repair 5, S5–S5 (2012).

28. Hinz, B. et al. The Myofibroblast One Function, Multiple Origins. Am J Pathology 170, 1807–1816 (2007).

29. Shook, B. A. et al. Myofibroblast proliferation and heterogeneity are supported by macrophages during skin repair. Science 362, eaar2971 (2018).

30. Mascharak, S. et al. Preventing Engrailed-1 activation in fibroblasts yields wound regeneration without scarring. Science 372, eaba2374 (2021).

31. Correa-Gallegos, D. et al. Patch repair of deep wounds by mobilized fascia. Nature 576, 287–292 (2019).

32. Shook, B. A. et al. Dermal Adipocyte Lipolysis and Myofibroblast Conversion Are Required for Efficient Skin Repair. Cell Stem Cell 26, 880–895.e6 (2020).

33. Guerrero-Juarez, C. F. et al. Single-cell analysis reveals fibroblast heterogeneity and myeloid-derived adipocyte progenitors in murine skin wounds. Nat Commun 10, 650 (2019).

34. Currie, J. D. et al. Live Imaging of Axolotl Digit Regeneration Reveals Spatiotemporal Choreography of Diverse Connective Tissue Progenitor Pools. Dev Cell 39, 411–423 (2016).

35. Guimarães-Camboa, N. et al. Pericytes of Multiple Organs Do Not Behave as Mesenchymal Stem Cells In Vivo. Cell Stem Cell 20, 345–359.e5 (2017).

36. Parfejevs, V., Antunes, A. T. & Sommer, L. Injury and stress responses of adult neural crest-derived cells. Dev Biol 444, S356–S365 (2018).

37. Chen, J. et al. Gli1+ Cells Couple with Type H Vessels and Are Required for Type H Vessel Formation. Stem Cell Rep 15, 110–124 (2020).

38. Tomasek, J. J., Gabbiani, G., Hinz, B., Chaponnier, C. & Brown, R. A. Myofibroblasts and mechano-regulation of connective tissue remodelling. Nat Rev Mol Cell Bio 3, 349–363 (2002).

39. Shin, D. & Minn, K. W. The Effect of Myofibroblast on Contracture of Hypertrophic Scar. Plast Reconstr Surg 113, 633–640 (2004).

40. Maurer, M., Peters, E. M. J., Botchkarev, V. A. & Paus, R. Intact hair follicle innervation is not essential for anagen induction and development. Arch Dermatol Res 290, 574–578 (1998).

41. Driskell, R. R., Giangreco, A., Jensen, K. B., Mulder, K. W. & Watt, F. M. Sox2-positive dermal papilla cells specify hair follicle type in mammalian epidermis. Development 136, 2815–2823 (2009).

42. Bai, C. B., Auerbach, W., Lee, J. S., Stephen, D. & Joyner, A. L. Gli2, but not Gli1, is required for initial Shh signaling and ectopic activation of the Shh pathway. Dev Camb Engl 129, 4753–61 (2002).

43. Madisen, L. et al. A robust and high-throughput Cre reporting and characterization system for the whole mouse brain. Nat Neurosci 13, 133–140 (2010).

44. Buch, T. et al. A Cre-inducible diphtheria toxin receptor mediates cell lineage ablation after toxin administration. Nat Methods 2, 419–426 (2005).

45. Schindelin, J. et al. Fiji: an open-source platform for biological-image analysis. Nat Methods 9, 676–682 (2012).

46. Rueden, C. T. et al. ImageJ2: ImageJ for the next generation of scientific image data. Bmc Bioinformatics 18, 529 (2017).

47. McQuin, C. et al. CellProfiler 3.0: Next-generation image processing for biology. Plos Biol 16, e2005970 (2018).

48. Bankhead, P. et al. QuPath: Open source software for digital pathology image analysis. Sci Rep-uk 7, 16878 (2017).

49. Pedregosa, F. et al. Scikit-learn: Machine Learning in Python. Arxiv (2012).

